# The CDK12 inhibitor SR-4835 functions as a molecular glue that promotes cyclin K degradation in melanoma

**DOI:** 10.1101/2023.05.30.542844

**Authors:** Thibault Houles, Jonathan Boucher, Geneviève Lavoie, Graham MacLeod, Sichun Lin, Stephane Angers, Philippe P. Roux

## Abstract

CDK12 is a transcriptional cyclin-dependent kinase (CDK) that interacts with cyclin K to regulate different aspects of gene expression. The CDK12-cyclin K complex phosphorylates several substrates, including RNA polymerase II (Pol II), and thereby regulates transcription elongation, RNA splicing, as well as cleavage and polyadenylation. Because of its implication in cancer, including breast cancer and melanoma, multiple pharmacological inhibitors of CDK12 have been identified to date, including THZ531 and SR-4835. While both CDK12 inhibitors affect Poll II phosphorylation, we found that SR-4835 uniquely promotes cyclin K degradation via the proteasome. Using loss-of-function genetic screening, we found that SR-4835 cytotoxicity depends on a functional CUL4-RBX1-DDB1 ubiquitin ligase complex. Consistent with this, we show that DDB1 is required for cyclin K degradation, and that SR-4835 promotes DDB1 interaction with the CDK12-cyclin K complex. Docking studies and structure-activity relationship analyses of SR-4835 revealed the importance of the benzimidazole side-chain in molecular glue activity. Together, our results indicate that SR-4835 acts as a molecular glue that recruits the CDK12-cyclin K complex to the CUL4-RBX1-DDB1 ubiquitin ligase complex to target cyclin K for degradation.

## INTRODUCTION

CDK12 (CRK7, CrkRS), along with its paralog CDK13 (CDC2L5, CHED), are amongst several cyclin-dependent kinases (CDKs) that regulate transcription ^1^. Specifically, CDK12 interacts with cyclin K to phosphorylate RNA polymerase II (Pol II) within its C-terminal domain (CTD), which serves as a platform for controlling transcriptional and post-transcriptional events ^2, 3^. Pol II phosphorylation by CDK12 regulates transcription elongation, splicing, as well as cleavage and polyadenylation ^4^. As CDK12 promotes elongation, it reduces the likelihood of cleavage at internal polyadenylation sites and increases the probability of cleavage at 3’ polyadenylation sites ^5, 6^. Several DNA damage response (DDR) genes have multiple intronic polyadenylation sites and were thus shown to be particularly sensitive to CDK12 inhibition ^5^. For this reason, CDK12 is thought to play important roles in genomic stability ^2, 7^, which underscores its potential as a therapeutic target.

Increasing evidence demonstrate the involvement of CDK12 in cancer ^8, 9^. In HER2- positive breast cancer, *CDK12* was shown to be co-amplified with the *HER2* oncogene, and to promote migration and invasion of HER2-positive breast tumor cells ^10^. CDK12 plays an important role in the growth of cancers driven by dysregulated transcription factors, such as MYC (neuroblastoma) ^11^ and the EWS–FLI1 fusion (Ewing sarcoma) ^12^, or in melanoma driven by activating mutations in *BRAF* ^13^. In other cancers, including ovarian ^14^, prostate ^15^, and breast ^16^, loss of *CDK12* leads to the downregulation of DDR genes, which increases genomic instability and sensitivity to chemotherapy. CDK12 can thus be used as biomarker, but also as a potential therapeutic target.

Several CDK12 inhibitors have been shown to have potential for cancer therapy. THZ1 is an irreversible covalent inhibitor of CDK7 that has therapeutic effects in several types of cancer^17^. At high concentrations, THZ1 can also inhibit CDK12, which was exploited in the design of THZ531 ^18^. THZ531 inhibits both CDK12 and CDK13, and studies have shown that THZ531 reduces Pol II phosphorylation and the proliferation of many types of cancer cells ^12, 13, 18, 19^. SR-4835 is another highly selective CDK12/13 inhibitor that act as an ATP antagonist ^16^. SR-4835 was shown to reduce Pol II phosphorylation and to induce rapid tumor regression in multiple mouse models ^16^. Recently, three groups identified molecules that bind to the CDK12-cyclin K complex and act as molecular glues to stabilize an interaction with the CUL4-RBX1-DDB1 ubiquitin ligase complex ^20–22^. These compounds were shown to promote cyclin K ubiquitination without the need for a DCAF substrate receptor, and to promote its degradation by the proteasome. These discoveries suggested that other CDK12/13 inhibitors may have hidden degrader activity ^23^.

Herein, we found that SR-4835 is another molecular glue-type cyclin K degrader. Our results show that the CUL4-RBX-DDB1 ubiquitin ligase complex is required for this effect, and structure-activity (SAR) relationship analysis suggests the importance of the benzimidazole side-chain in molecular glue activity.

## RESULTS

### Characterization of SR-4835 in BRAF-mutated melanoma cells

BRAF-mutated melanoma cells were recently shown to display high CDK12 activity ^13^. To assess its role in these cells, we treated five BRAF-mutated melanoma cell lines (Colo829, A375, WM164, WM983A, WM983B) with increasing concentrations of SR-4835, a recently described ATP-competitive inhibitor of CDK12 and CDK13 ^16^. We found that SR-4835 inhibited the proliferation of all cell lines with IC_50_ values ranging from 80.7 to 160.5 nM (Fig. 1A). To assess CDK12 activity, we determined the impact of SR-4835 on the phosphorylation of RPB1 (Fig. 1B), a Pol II subunit and specific substrate of CDK12 ^24^. We found that SR-4835 treatment (6 hrs) strongly decreased RPB1 phosphorylation at Ser2, but not at Ser5, which is indicative of reduced CDK12/13 activity ^25^. As CDK12 inhibition was previously shown to decrease the expression of many DDR genes ^2^, we quantified the abundance of *BRCA1*, *RAD51*, *ATM*, *XRCC2*, *FANCI* and *FANCD2* in response to two different doses of SR-4835 (30 and 100 nM). As expected, SR-4835 treatment (6 hrs) decreased the expression of all DDR genes tested (Fig. 1C), which was also seen at the protein level for BRCA1, ATM and RAD51 when treatment was extended for 24 hours (Fig. 1D). As expected, these data correlated with the accumulation of DNA damage in response to SR-4835 treatment, which was shown using γH2AX phosphorylation (Fig. 1D). Together, these results indicate that SR-4835 efficiently inhibits CDK12/13 activity in BRAF-mutated melanoma, which results in increased DNA damage and reduced cell proliferation.

**Fig. 1:**
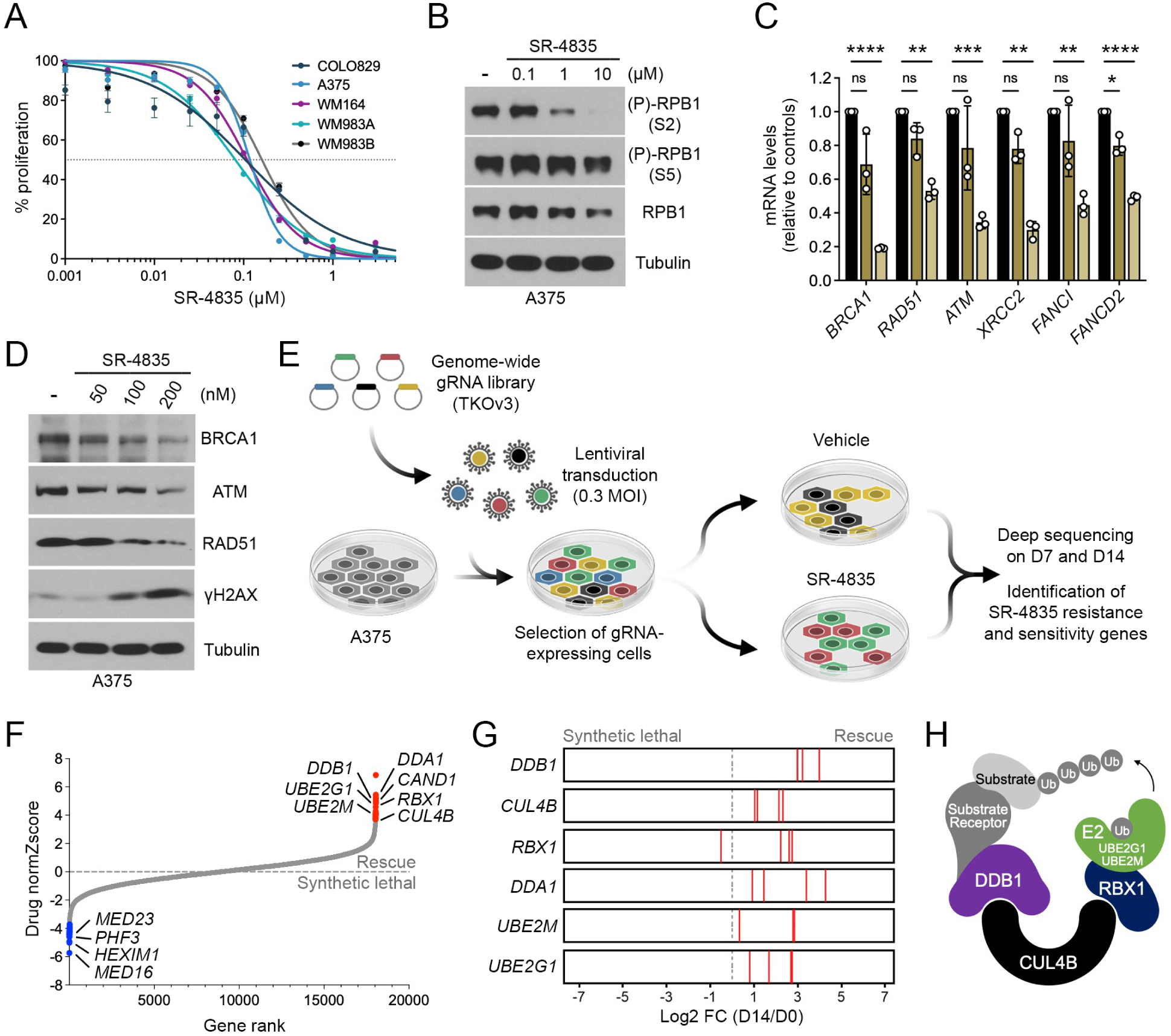
Characterization of SR-4835 in BRAF-mutated melanoma cells. **A** Proliferation assay was performed using a panel of BRAF-mutated melanoma cell lines treated with increasing doses of SR-4835 for 96 h. Respectively, the IC_50_ for A375, 117.5 nM; Colo829, 104.5 nM; WM164, 109.6 nM; WM983A, 80.7 nM; WM983B 160.5 nM. **B** Immunoblot depicting RPB1 phosphorylation levels in A375 cells treated with SR-4835 (0.1, 1 and 10 µM) for 6 h. **C** qPCR analysis from A375 cells treated for 6 h with SR-4835 (30 and 100 nM). **D** Immunoblot of A375 cells treated for 24 h with SR-4835 (0.05, 0.1 and 0.2 µM). **E** Schematic representation of the CRISPR/Cas9 screen used to characterize SR-4835 activity. **F** Waterfall depicting differential gene essentiality in A375 treated with SR-4835 based on comparison of gRNA abundance before and after 14 days of treatment. **G** Bar graphs illustrating the Log2 fold-change of the four gRNAs targeting DDA1, DDB1, UBE2M, UBE2G1, CUL4B, RBX1 genes at day 14 in A375 treated with SR-4835. **H** Schematic representation of the CRL4B complex (RBX1-CUL4B- DDB1). (**A**, **C**) Data are represented as mean ± SD of independent experiment, n = 3. Representative data of n = 3 (**B**, **D**) or normZscore of independent sgRNA, n = 4 (**F**). Significance was determined using unpaired two-tailed Student’s t-tests (**C**). *P*-values signification: ns, not significant; * *P* < 0.05; ** *P* < 0.01; *** *P* < 0.01; **** *P* < 0.001.

To characterize the mechanism of action of SR-4835, we performed a loss-of-function CRISPR/Cas9 screen in A375 cells infected with a genome-wide lentiviral gRNA library (Fig. 1E). For this, 18 053 genes were targeted using 70 948 gRNAs (4 guides/gene) in cells subjected to vehicle (DMSO) or SR-4835 treatment (50 nM). Cells were allowed to grow for seven (D7) and fourteen (D14) days before analyzing gRNA abundance respective to control cells (D0). Essentiality scores were calculated using DrugZ ^26^, which revealed 19 synthetic lethal genes (blue) and 19 rescue genes (red) (Fig. 1F; Dataset 1). We found significant enrichments in specific GO terms, including “RNA Pol II transcription regulator complex” (Log(p-value) = -3.64) amongst synthetic lethal genes, and “Cul4B-RING E3 ubiquitin ligase complex” (Log(p-value)= -8.68) amongst rescue genes. Specifically, rescue genes appeared to comprise genes coding for five members of a CRL4B complex (DDB1, CUL4B, RBX1, DDA1), and two genes coding for E2 ubiquitin-conjugating enzymes (UBE2G1 and UBE2M) (Fig. 1F). These data were confirmed by analyzing the impact of each individual gRNAs (Fig. 1G). Taken together, these results suggest that SR-4835 requires a functional CUL4-RBX1-DDB1 ubiquitin ligase complex to inhibit the proliferation of BRAF-mutated melanoma cells (Fig. 1H).

### SR-4835 promotes the proteasomal degradation of cyclin K

The CUL4-RBX1-DDB1 ubiquitin ligase complex was previously shown to promote cyclin K degradation in response to CDK12 inhibitors, including CR8, HQ461 and dCeMM2/3/4 ^20–22^. To determine if SR-4835 promotes cyclin K degradation via similar mechanisms, A375 cells were treated with increasing concentrations of SR-4835 (2 hrs) and assessed for cyclin K levels. Compared to THZ531, which is a covalent CDK12/13 inhibitor that does not promote cyclin K degradation ^20–22^, we found that SR-4835 reduced cyclin K levels in a dose-dependent manner (Fig. 2A). This effect was not specific to a particular cell line, as we found that SR-4835 treatment (2 hrs) of three additional BRAF-mutated melanoma cell lines (WM164, WM983B and COLO829) also resulted in reduced cyclin K levels (Fig. 2B). To determine if SR-4835 affects cyclin K stability, A375 cells were treated with DMSO or SR-4835 (1 µM) over a time-course of cycloheximide (100 µg/mL) treatments (0 to 120 min) to block *de novo* protein synthesis (Fig. 2C). In DMSO-treated A375 cells, we found that cyclin K is a relatively stable protein with an apparent half-life that exceeds 120 min (Fig. 2C, 2D). However, cyclin K protein levels were found to rapidly decrease in the presence of SR-4835, resulting in an apparent half-life of 47.8 min (Fig. 2C, 2D). This effect appeared to occur at the protein level, as SR-4835 treatment (1 µM) did not affect *CCNK* mRNA levels (Fig. 2E).

**Fig. 2:**
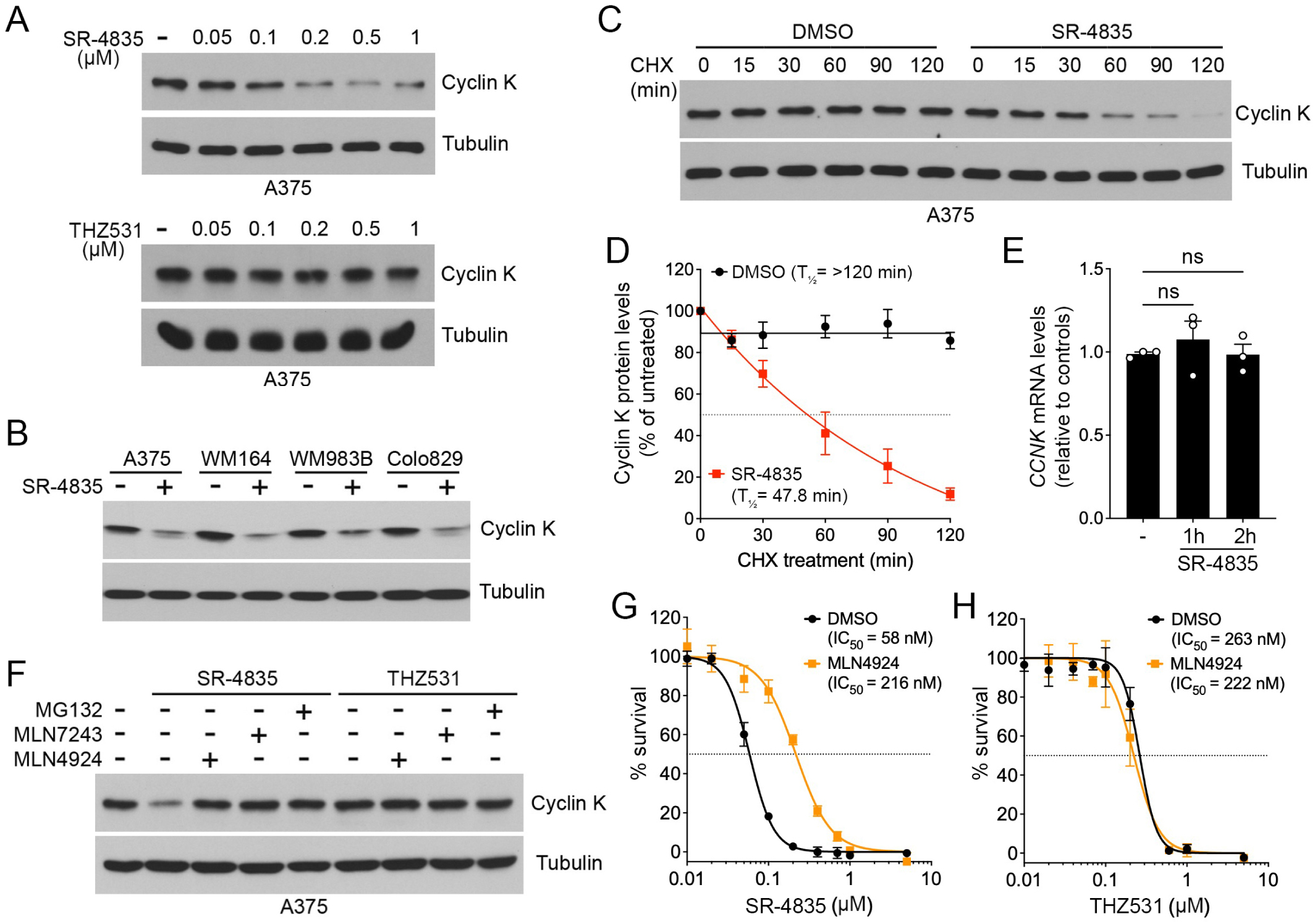
SR-4835 promotes the proteasomal degradation of cyclin K. **A** Immunoblot depicting cyclin K levels in A375 cells treated with SR-4835 or THZ531 (0.05, 0.1, 0.2, 0.5 and 1 µM) for 2 h. **B** Immunoblot depicting cyclin K levels in melanoma cell lines treated with SR-4835 (1 µM) for 2 h. **C** A375 cells were treated with vehicle (DMSO) or SR-4835 (1 µM) during a time course of cycloheximide (CHX) treatment (100 µg/mL). Endogenous cyclin K levels was analyzed by immunoblotting. **D** Densitometric analysis of cyclin K levels was performed on CHX time course shown in **C**. The data were normalized to tubulin and then expressed relative to respective controls (t = 0). **E** qPCR of A375 cells treated with SR-4835 (1 µM) for indicated times. **F** A375 cells were treated with MG132 (10 µM), MLN7243 (1 µM) or MLN4924 (0.5 µM) for 1h before 2 h of SR-4835 or THZ531 treatment (1 µM). **G** Proliferation assay was performed in A375 cells treated with SR-4835 at indicated doses combined with MLN4924 (0.5 µM) for 72 h. Respectively, IC_50_ value for SR-4835 alone is 58 nM, and 216 nM when in combination with MLN4924. **H** Same as **G**, except that A375 cells were treated with THZ531. IC_50_ concentration for A375 with THZ531 alone is 263 nM, and 222 nM when combined with MLN4924. (**A**, **B**, **F** - **H**) Representative data of n = 3 or n = 4 (**C**). Data are represented as mean ± SD of independent experiments, n = 4 (**D**), n = 3 (**E**) or independent replicate, n = 4 (**G**, **H**). Significance was determined using unpaired two-tailed Student’s t-tests (**E**). *P*-values signification: ns, not significant.

To determine if cyclin K depletion requires the ubiquitin-proteasome system (UPS), A375 cells were treated with inhibitors of the E1 ubiquitin-activating enzyme (MLN7243), the NEDD8- activating enzyme (MLN4924) and the proteasome (MG132), prior to CDK12 inhibition. While SR-4835 treatment alone reduced cyclin K levels, we found that addition of all three inhibitors prevented this effect, suggesting that SR-4835 promotes cyclin K degradation via the UPS (Fig. 2F). Conversely, none of the inhibitors affected cyclin K protein levels in the presence of THZ531, indicating that this effect was specific to SR-4835. MLN4924 efficiently blocks neddylation of all cullins, leading to the accumulation of their substrates, such as cyclin K. To determine if MLN4924 treatment could reduce SR-4835 cytotoxicity, A375 cells were treated with DMSO or MLN4924 (0.5 µM) and increasing concentrations of SR-4835 or THZ531 for 72 hrs. Consistent with its effect on cyclin K, we found that MLN4924 prevented SR-4835 cytotoxicity by 3.7-fold (Fig. 2G). This effect was not seen with THZ531 (Fig. 2H), which does not promote cyclin K depletion (Fig. 2A). Together, these results suggest that cullin neddylation and cyclin K degradation substantially contribute to SR-4835 cytotoxicity.

### Active-site engagement of SR-4835 promotes CDK12 binding to DDB1

Based on the identification of *DDB1* as a knockout gene that rescues from SR-4835 cytotoxicity (Fig. 1F), we assessed whether cyclin K degradation requires the CUL4 adaptor protein DDB1. To verify this, we depleted DDB1 from A375 cells using two lentiviral vectors expressing different short hairpin RNA (shRNA) (Fig. 3A). We found that shRNA-mediated DDB1 knockdown conferred resistance to cyclin K degradation by SR-4835 (Fig. 3A), suggesting the implication of a CUL4- RBX1-DDB1 ubiquitin ligase complex. Several small-molecules that promote cyclin K degradation were reported to act as molecular glues between CDK12 and DDB1 ^20–22^. To determine if SR-4835 also promotes the interaction between CDK12 and DDB1, we performed co-immunoprecipitation experiments in HEK293 cells transfected with Myc-CDK12, Flag-DDB1 and HA-cyclin K. In the absence of inhibitors, we found a basal interaction between CDK12 and DDB1 (Fig. 3B), suggesting that CDK12 and DDB1 may naturally interact in cells. This interaction increased in response to SR-4835 treatment in a time-dependent manner, which correlated with cyclin K degradation (Fig. 3B). The interaction between CDK12 and DDB1 was not increased by THZ531 treatment, consistent with the fact that this inhibitor does not promote cyclin K degradation (Fig. 2A). As SR-4835 and THZ531 are both ATP-competitive CDK12 inhibitors, we determined whether covalent binding of THZ531 to the active site of CDK12 may prevent SR-4835 from promoting cyclin K degradation. For this, A375 cells were pretreated with DMSO or different concentrations of THZ531 (2 hrs) and exposed to SR-4835 (1 µM) for 2 hrs to promote cyclin K degradation. We found that THZ531 treatment efficiently prevented cyclin K degradation induced by SR-4835 (Fig. 3C), suggesting that saturating the ATP-binding pocket of CDK12 prevents SR-4835 binding and thereby rescues cyclin K destabilization. Molecular modeling studies using the X-ray structure of CDK12 in association with DDB1 and cyclin K predicts the binding mode of SR-4835 to be ATP-competitive (Fig. 3D) and pointed towards DDB1 (Fig. 3E), as was similarly shown for CR-8 ^20^. Together, these data suggests that CDK12 active-site engagement is required for the cyclin K destabilizing effect of SR-4835.

**Fig. 3:**
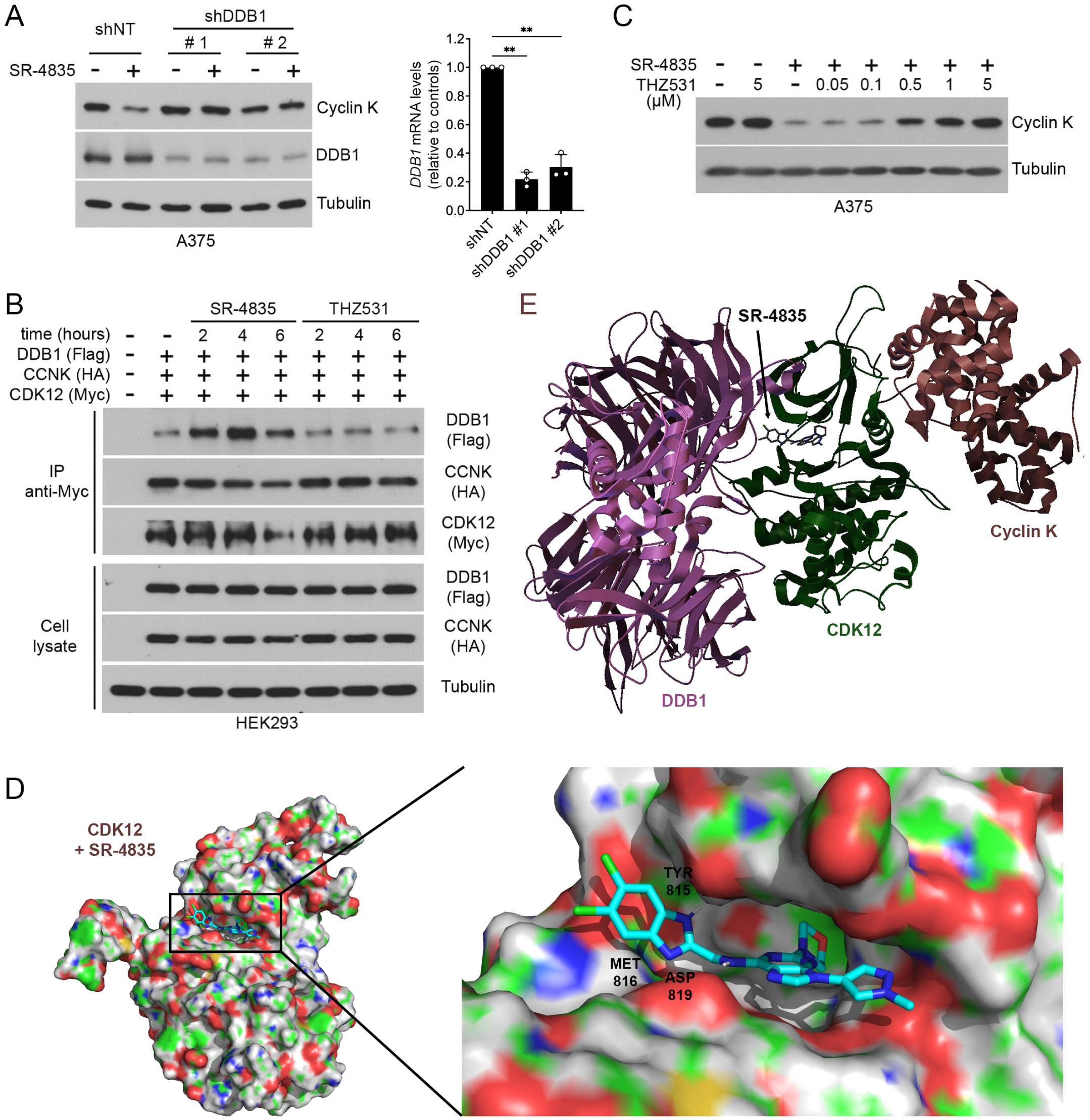
Active-site engagement of SR-4835 promotes CDK12 binding to DDB1. **A** Immunoblot depicting cyclin K levels in A375 cells depleted for DDB1 using shRNA constructs and treated with SR-4835 (1 µM) for 2 h. shRNA efficiency was assessed by immunoblotting with the anti-DDB1 antibody, or by qPCR. **B** HEK293 cells were transfected with Myc-tagged CDK12, HA-tagged cyclin K and Flag-tagged DDB1, and treated for indicated times with SR- 4835 (1 µM) or THZ531 (1 µM). Immunoprecipitated Myc-CDK12 was assayed for co-IP with DDB1 and cyclin K. **C** A375 cells were treated with THZ531 at indicated concentrations for 45 min before SR-4835 (1 µM) treatment for another 2 h. Cyclin K levels were assessed by immunoblot. **D** Docking pose of SR-4835 in the active site of CDK12 (PDB: 6TD3) with key residues highlighted. **E** Docking pose of SR-4835 in the active site of CDK12 in association with DDB1 and cyclin K (PDB: 6TD3). (**A**, **B**, **C**) Representative data of n = 3. Data are represented as mean ± SD of independent experiments (n = 3) and significance was determined using unpaired two-tailed Student’s t-tests (**A**). *P*-values signification: ** *P* < 0.01.

### The benzimidazole side-chain of SR-4835 is important for cyclin K degradation and cytotoxicity

To characterize the molecular glue activity of SR-4835, we conducted a SAR study. Based on its resemblance to CR8 (Fig. 4A), which binds to the active site of CDK12 and uses a surface-exposed phenylpyridine ring to interact with DDB1 ^20^, we targeted the similarly positioned dichlorobenzimidazole of SR-4835. As described elsewhere ^27^, we generated structural analogs that probed the contribution of the benzimidazole side-chain, which included the corresponding 5,6-difluorobenzathiazole (CU-0904), 4,5-difluorobenzathiazole (CU-0905), and 5-methoxy,6-fluorobenzathiazole (CU-0906) (Fig. 4A). To evaluate their activity, we first measured cyclin K abundance in A375 cells treated with increasing concentrations of each compound (2 hrs). While we found that all analogues could promote cyclin K degradation to different extents (EC_50_ values between 197 and 333 nM), their activity remained lower than that of SR-4835 (EC_50_ of 112 nM) (Fig. 4B).

**Fig. 4:**
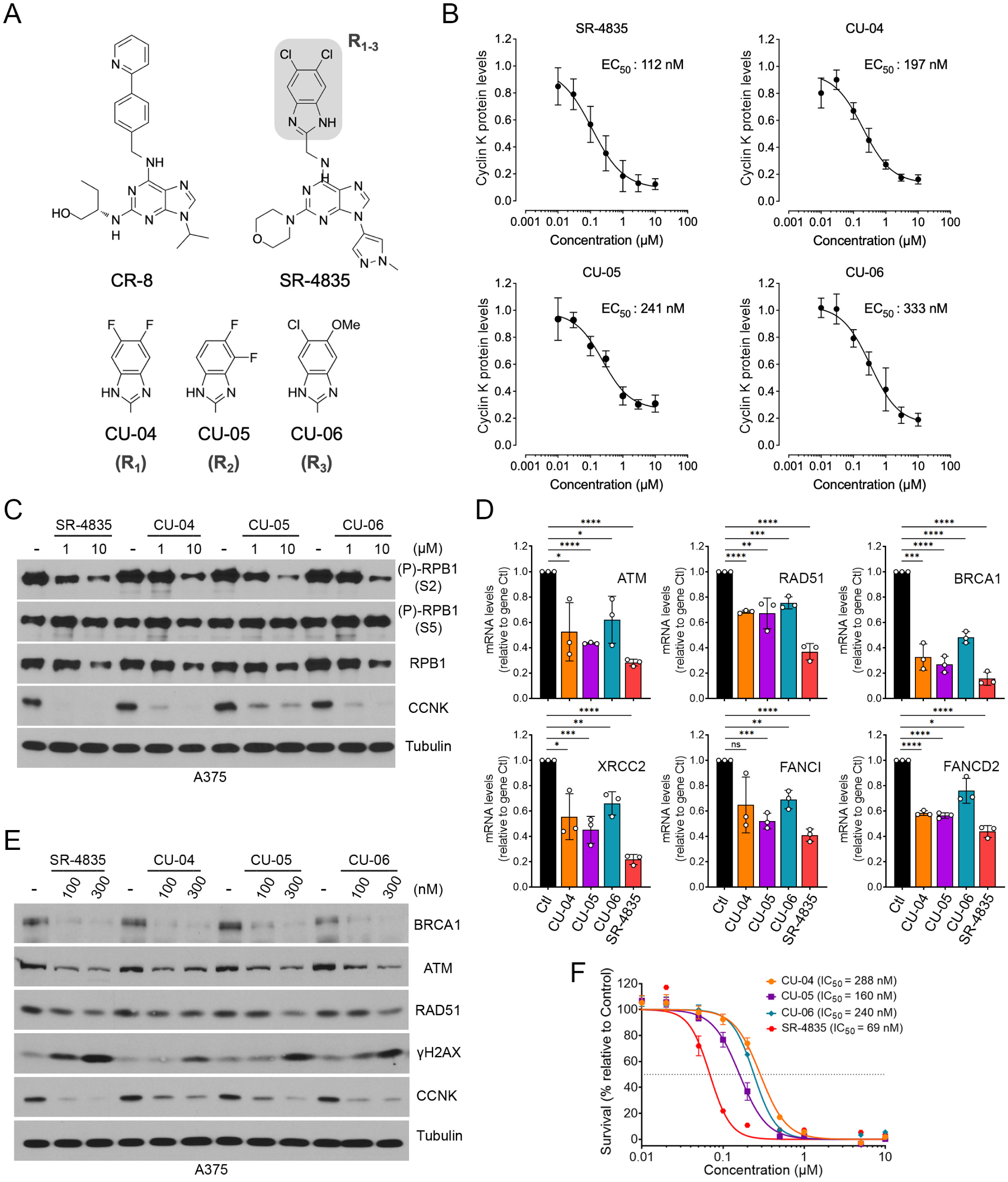
The benzimidazole side-chain of SR-4835 modulates its molecular glue activity. **A** Chemical structure of CR8 and SR-4835, alongside developed analogues. **B** A375 cells were treated with SR-4835, CU-0904, CU-0905 or CU-0906 for 2 h at indicated concentrations, and cyclin K levels were quantified by densitometric analysis and normalized to tubulin. EC_50_ concentration for SR-4835 was 112 nM, 197 nM for CU-0904, 241 nM for CU-0905 and 333 nM for CU-0906. **C** Immunoblot depicting RPB1 phosphorylation levels in A375 cells treated with SR-4835, CU-0904, CU-0905 or CU-0906 (1 and 10 µM) for 6 h. **D** qPCR of A375 cells treated for 6 h with SR-4835, CU-0904, CU-0905 or CU-0906 (100 nM). **E** Immunoblot of A375 cells treated with SR-4835, CU-0904, CU-0905 or CU-0906 (100 and 300 nM) for 24 h. **F** Proliferation assay was performed using A375 cells treated with increasing doses of SR-4835, CU-0904, CU-0905 or CU-0906 for 72 h. IC_50_ concentration for A375 treated with SR-4835 was 69 nM, 288 nM for CU-0904, 160 nM for CU-0905 and 240 nM for CU-0906. (**B**) Data are represented as mean ± SEM of independent experiments, n = 3. (**C**, **E**, **F**) Representative data of n = 3. Data are represented as mean ± SD of independent experiment, n = 3 (**D**) or independent replicate, n = 6 (**F**). *P*-values signification: ns, not significant; * *P* < 0.05; ** *P* < 0.01; *** *P* < 0.01; **** *P* < 0.001.

We also tested the analogues for their ability to inhibit CDK12 and thereby reduce RPB1 phosphorylation at Ser2. We found that all analogues reduced RPB1 phosphorylation in a dose-dependent manner, but that SR-4835 appeared to be more potent (Fig. 4C). We also found that all drugs could inhibit DDR gene expression (*ATM*, *RAD51* and *BRCA1*), as evaluated by qPCR (Fig. 4D) and by immunoblotting for protein levels (Fig. 4E). In both cases, SR-4835 appeared more potent, which also correlated with the accumulation of DNA damage, as shown using γH2AX phosphorylation (Fig. 4E). Finally, we determined the impact of the different analogues on A375 cytotoxicity. We found that CU-0904, CU-0905 and CU-0906 reduced cell proliferation with similar IC_50_, consisting of 288, 160 and 240 nM, respectively (Fig. 5F). Here again, SR- 4835 was more potent than its analogues to inhibit cell proliferation, with an IC_50_ of 69 nM. These experiments characterized the molecular glue activity of SR-4835 analogs, and confirmed the role played by the benzimidazole side-chain for cyclin K degradation.

## DISCUSSION

CDK12 is an exploitable vulnerability in multiple types of cancer ^8, 23^, including melanoma ^13^, making it an attractive therapeutic target. To characterize the mechanism of action of SR-4835, a recently described CDK12 inhibitor ^16^, we performed a genome-wide CRISPR/Cas9 loss-of-function screen in A375 melanoma cells. We found that major components of the CRL4B E3 ubiquitin ligase complex, including DDB1, CUL4B, RBX1, and DDA1, along with E2 ubiquitin-conjugating enzyme subunits, rescued cells from SR-4835 cytotoxicity (Fig. 1). Furthermore, we found that SR-4835 shortens the half-life of cyclin K through the UPS (Fig. 2), leading us to hypothesize that SR-4835 directs the CRL4B complex to the CDK12-cyclin K heterodimer. This function, known as molecular glue degrader, has been observed in other CDK12 inhibitors such as dCEMM2-4, HQ461, CR8, and NCT02 ^20–22, 28^. Here, we demonstrate that SR-4835-induced cyclin K degradation depends on the adaptor protein DDB1 and occurs through increased DDB1-CDK12 physical interaction (Fig. 3). SAR analysis of SR-4835 revealed the importance of the benzimidazole side-chain in molecular glue activity and resulted in SR-4835 derivates with varying potency (Fig. 4). Together, our results show that SR-4835 functions as a molecular glue between the CDK12-cyclin K heterodimer and the CUL4B complex. Structural analysis will be required to better define the roles of each of CRL4B complex components in cyclin K degradation induced by SR-4835.

Canonical molecular glue degraders typically require DDB1- and CUL4-associated factors (DCAFs) to promote DDB1 association with its targeted substrate ^29^. Previous studies have also shown that cyclin K degraders can non-canonically bypass the requirement for DCAF by directly interacting with DDB1 ^20, 21^. Interestingly, our CRISPR/Cas9 screen identified several DCAFs that either rescued or sensitized cells to SR-4835 cytotoxicity, including WDR82 and WDR12, or DCAF8, DCAF16, DCAF4L1 and TLE1, respectively (Dataset 1). These results suggest that some DCAFs can either compete or favor SR-4835 cytotoxicity, presumably by modulating the free pool of DDB1 in cells. However, it remains an option question whether these DCAFs can affect SR-4835-mediated cyclin K degradation.

Once bound to the ATP-binding pocket of CDK12, molecular glues such as CR8 expose critical domains that interact with specific residues within the BPC of DDB1 ^20^. The structure of surface-exposed moieties determines the stability of the DDB1-CDK12 complex and ultimately affects the degrader activity of molecular glues. For CR8, a cyclin K molecular glue closely related to SR-4835, the surface-exposed hydrophobic phenylpyridine system, which is made up of 2-pyridyl moieties, was shown to mediate its optimal efficacy ^20^. Roscovitine, the parent compound of CR8 that lacks one of the two pyridyl groups, promoted weaker complex formation between CDK12 and DDB1 and failed in inducing cyclin K degradation. These observations strongly support the contribution of this CR8-related sidechain for optimal CRL4-dependent molecular glue activity. Given its resemblance to CR8, we tested the importance of the related dichlorobenzimidazole sidechain in SR-4835. Indeed, docking studies indicated that SR-4835 is similarly positioned in the active site of CDK12, with its dichlorobenzimidazole sidechain pointing towards DDB1 (Fig. 3E). We generated three structural analogs with varying chemical groups attached to its benzimidazole sidechain, as previously reported ^27^. We found that all these analogs exerted CDK12 inhibitory activity and promoted cyclin K degradation to different capacities, but to a lesser extent than SR-4835. These results confirmed the importance of the benzimidazole moiety and justify the relevance of additional studies to further optimize the potency of SR-4835 as a molecular glue degrader.

## METHODS

### CRISPR/Cas9-based screening

1.6 x 10^8^ A375 cells were transduced into thirty-two 150 mm plates with lentivirus carrying The Toronto Knockout Library v3 (TKOv3, Addgene # 90294) in standard culture media + 8 µg/mL polybrene. Lentiviral transduction was performed at a multiplicity of infection of ∼ 0.3 to ensure that most cells would contain a single lentiviral insertion following selection. Twenty-four hours post-transduction, viral media was removed, and cells were subjected to 2 µg/mL puromycin selection for 2 days. After selection, transduced cells were trypsinized and D0 samples of 20 million cells were collected for genomic DNA extraction. The remaining cells were split into 4 replicates of 20 million cells. Two replicates were treated with 50 nM SR-4835 and the remaining two treated with an equivalent volume of vehicle (DMSO). Cells were cultured for 14 days in total with regular media changes and passaging, with a minimum of 20 million cells per replicate at all times to maintain a library coverage of greater than 200-fold. Cell pellets were collected for genomic DNA extraction at day 7 (D7) and day 14 (D14).

Genomic DNA was extracted from D0, D7 and D14 cell pellets using the QIAamp DNA Blood Maxi Kit (QIAGEN) according to the manufacturer’s protocol. 50 μg of gDNA from each sample was amplified in 20 individual PCR reactions using Kapa HiFi Master Mix (Kapa Biosystems) using TKOv3 library primers (forward: 5’-GAGGGCCTATTTCCCATGATTC-3’, reverse: 5’- GTTGCGAAAAAGAACGTTCACGG-3’) for 19 amplification cycles. Replicate PCR reactions for individual samples were pooled and 5 μL was used as a template for a second PCR reaction to attach Illumina sequencing adaptors and barcodes for each sample. Barcoded PCR products were gel purified and sequenced on an Illumina NextSeq500 instrument.

DNA sequencing reads were mapped to TKOv3 library gRNA sequences using MAGeCK version 0.5.8 ^30^. Read count data was then analyzed using the DrugZ python package to identify synergistic and suppressor gene-drug interactions ^26^.

### Cell culture

HEK293, A375, WM164 cell lines were purchased from ATCC. WM983A and WM983B cell lines were purchased from Rockland (Philadelphia, PA). All cell lines were maintained in Dulbecco’s modified Eagle’s medium (DMEM) (GIBCO) with 4.5 g/liter glucose supplemented with 10% fetal bovine serum (FBS), 100 IU/mL penicillin, and 100 µg/mL streptomycin, under 37°C/5%CO_2_ conditions. Colo829 cells were purchased from ATCC and maintained in Roswell Park Memorial Institute Medium (RPMI) with 10% FBS, 100 IU/mL penicillin and 100 µg/mL streptomycin, under 37°C/5%CO_2_ conditions. Cells were regularly tested by PCR to exclude mycoplasma contamination.

### DNA constructs and chemicals

The original plasmid encoding human CDK12 (pLP-3xFLAG- CDK12) and HA-CCNK were obtained from Gregg B. Morin (University of British Columbia). CDK12 DNA construct was used as PCR template for generating 6Myc-tagged CDK12 in pcDNA3. pcDNA3-FLAG-DDB1 was obtained from Addgene (#19918). SR-4835 was purchased from MedChemExpress and CU-0904, CU-0905, CU-0906 were purchased from Omegachem. All compounds used in this study are >95% pure by HPLC analysis (See Supporting Information).

### Immunoprecipitations and immunoblotting

Cell lysates were prepared as previously described ^31^. Briefly, cells were washed three times with ice-cold phosphate-buffered saline (PBS) and lysed in RIPA or BLB buffer (10 mM K_3_PO_4_, 1 mM EDTA, 5 mM EGTA, 10 mM MgCl_2_, 50 mM β-glycerophosphate, 0.5% Nonidet P-40, 0.1% Brij 35, 0.1% deoxycholic acid), complemented with 1 mM sodium orthovanadate [Na_3_VO_4_], 1 mM phenylmethylsulfonyl fluoride, and a cOmplete protease inhibitor cocktail tablet (Roche). Lysates were centrifuged at 16,000 × g for 10 min at 4 °C, supernatants were collected and heated for 10 min at 95 °C in 4 x Laemmli buffer (5 × buffer is 60 mM Tris-HCl [pH 6.8], 25% glycerol, 2% SDS, 14.4 mM 2- mercaptoethanol, and 0.1% bromophenol blue). Alternatively, for immunoprecipitations, cells were lysed in BLB buffer and supernatants were incubated with the indicated antibodies for 2 h, followed by 1 h of incubation with protein A–Sepharose CL-4B beads (GE Healthcare). Immunoprecipitates were washed three times in lysis buffer, and beads were eluted and heated in 2x Laemmli buffer. Eluates and total cell lysates were subjected to 7.5 to 12% SDS-PAGE, and resolved proteins were transferred onto polyvinylidene difluoride (PVDF) membranes for immunoblotting. The data presented are representative of at least three independent experiments. Antibodies targeted against CDK12, cyclin K and DDB1 are from Bethyl Laboratories. Antibodies against, RPB1 (4H8), p-RPB1 (S2) (E1Z3G), p-RPB1 (S5) (D9N5I), p- H2AX (S139) (20E3), BRCA1, ATM (D2E2) and Rad51 (D4B10) were purchased from Cell Signaling Technologies. Antibodies against Tubulin (T5618), Myc (9E10), Flag (2EL-1B11) are from Millipore Sigma and HA (12CA5) from ThermoFisher. All secondary horseradish peroxidase (HRP)-conjugated antibodies used for immunoblotting were purchased from Chemicon.

### Survival assays

Cells were grown in medium supplemented with 10% FBS and relative number of viable cells was measured using WST-1 reagent from Millipore Sigma. WST-1 was added 2 h prior to absorbance measurement at 450 nm using a Tecan Infinite M200 PRO microplate reader. Cells were plated and after adhesion were treated with SR-4835, CU-0904, CU-0905, CU-0906 (0.01 to 10 µM) or THZ531 (0.005 to 5 µM) in addition or not to MLN4924 at 75 nM. Cells were grown for 72 h before addition of WST1. The results displayed represent the mean of n = 3, 4 or 6 ± standard deviation (SD).

### Viral Infections and transfection

For shRNA-mediated knockdown of DDB1, lentiviruses were produced using vectors from the Mission TRC shRNA library (Sigma-Aldrich) targeting DDB1 (#1 TRCN0000082855, #2 TRCN0000082857). One day after infection, A375 cells were selected with 2 μg/ml puromycin. HEK293 and HEK293T cells were transfected by calcium phosphate precipitation. Briefly, plasmid and CaCl_2_ 2 M were diluted v/v in 2x HBS (274 mM NaCl, 1.5 mM Na_2_HPO_4_, 55 mM HEPES, pH 7) before adding slowly to cells.

### RNA extraction and real-time quantitative-PCR (qPCR)

Total RNA was extracted using RNeasy mini Kit (Qiagen) and reverse-transcribed using the cDNA Reverse Transcription Kit (Applied Biosystems), as described by the manufacturer. For each qPCR assay, a standard curve was performed to ensure the efficacy of the assay (between 90 and 110%). The Viia7 qPCR instrument (Life Technologies) was used to detect amplification level and was programmed with an initial step of 20 s at 95 °C, followed by 40 cycles of 1 s at 95 °C and 20 s at 60 °C. Relative expression (RQ = 2−ΔΔCT) was calculated using the Expression Suite software (Life Technologies), and normalization was done using both *ACTB* and *GAPDH*.

### Docking studies

Docking was performed using Autodock Vina, version 4.2 ^32^. The crystal structure of CDK12 (PDB ID: 6TD3) and the ligand SR-4835 were prepared using standard procedures and AutoDockTools 1.5.7. The grid box was defined to encompass the active site of CDK12 (center_x: -61.361, center_y: 21.611, center_z: -11.917, size_x: 52, size_y: 46, size_z: 56) and the scoring function was set to use default parameters.

### Quantification and statistical analysis

For all relevant panels unless specified, center values and error bars represent means ± SD and *p*-values were determined by unpaired two-tailed Student’s t-tests used for comparisons between two groups with at least n = 3. *P*-values signification: ns, not significant; *, *P* < 0.05; **, *P* < 0.01; ***, *P* < 0.001; ****, *P* < 0.0001. All experiments were performed a minimum of three times. Statistical tests for each experiment are detailed in the results, figures, and figure legends.

## ACKNOWLEDGMENTS

We thank all members of the laboratory for their insightful discussions and comments. We thank Dr. Gregg B. Morin (University of British Columbia) for providing DNA constructs. We are grateful for the expertise of personnel at the IRIC Genomics core facility. This work was supported by grants from the Cancer Research Society, the Canadian Institutes for Health Research, and from the Natural Sciences and Engineering Research Council of Canada (to P.P.R.).

